# The *TMEM106B* rs1990621 protective variant is also associated with increased neuronal proportion

**DOI:** 10.1101/583286

**Authors:** Zeran Li, Fabiana G. Farias, Umber Dube, Jorge L. Del-Aguila, Kathie A. Mihindukulasuriya, Maria Victoria Fernandez, Laura Ibanez, John P. Budde, Fengxian Wang, Allison M. Lake, Yuetiva Deming, James Perez, Chengran Yang, Joseph Bradley, Richard Davenport, Kristy Bergmann, Bruno A. Benitez, Joseph D. Dougherty, Oscar Harari, Carlos Cruchaga

## Abstract

**Background:** In previous studies, we observed decreased neuronal and increased astrocyte proportions in AD cases in parietal brain cortex by using a deconvolution method for bulk RNA-seq. These findings suggested that genetic risk factors associated with AD etiology have a specific effect in the cellular composition of AD brains. The goal of this study is to investigate if there are genetic determinants for brain cell compositions.

**Methods:** Using cell type composition inferred from transcriptome as a disease status proxy, we performed cell type association analysis to identify novel loci related to cellular population changes in disease cohort. We imputed and merged genotyping data from seven studies in total of 1,669 samples and derived major CNS cell type proportions from cortical RNAseq data. We also inferred RNA transcript integrity number (TIN) to account for RNA quality variances. The model we performed in the analysis was: normalized neuronal proportion ∼ SNP + Age + Gender + PC1 + PC2 + median TIN.

**Results:** A variant rs1990621 located in the *TMEM106B* gene region was significantly associated with neuronal proportion (p=6.40×10^−07^) and replicated in an independent dataset. The association became more significant as we combined both discovery and replication datasets in multi-tissue meta-analysis (p=9.42×10^−09^) and joint analysis (p=7.66×10^−10^). This variant is in high LD with rs1990622 (r^2^ = 0.98) which was previously identified as a protective variant in FTD cohorts. Further analyses indicated that this variant is associated with increased neuronal proportion in participants with neurodegenerative disorders, not only in AD cohort but also in cognitive normal elderly cohort. However, this effect was not observed in a younger schizophrenia cohort with a mean age of death < 65. The second most significant loci for neuron proportion was *APOE*, which suggested that using neuronal proportion as an informative endophenotype could help identify loci associated with neurodegeneration.

**Conclusion:** This result suggested a common pathway involving *TMEM106B* shared by aging groups in the present or absence of neurodegenerative pathology may contribute to cognitive preservation and neuronal protection.

## Introduction

Although neuronal loss and synapse dysfunction are the preceding events of cognitive deficits in Alzheimer’s disease (AD), neurons do not work or survive by themselves. These delicate organelles require supports through intimate collaborations within themselves and with other cell types^1^. The microenvironment of cellular crosstalk, interaction, balance, and circuits maintained by neurons, astrocytes, microglia, oligodendrocytes, and other vascular cells are essential for the brain to carry out functions and fight against insults.

AD associated risk factors identified across the genome also point to the involvements of multi-cell types apart from neurons^1,2^. APOE4 is related to lipid metabolism and mostly expressed in astrocyte and microglia^3^. Other lipid metabolism related risk genes are *ABCA7* identified in all cell types^2,4^, *CLU* in astrocyte and oligodendrocyte precursor cells^2,5,6^, and *SORL1* in astrocyte^2^. Research interests in the roles of inflammatory response to toxic stimuli or microbial infection have been escalating recently, and AD risk genes associated with immune response including *TREM2*^7^-^9^, *PLCG2*^9^, *ABI3*^9^, *CR1*^2,6^, *CD33*^2,4^, *HLA-DRB5–HLA-DRB1*^2^, and *INPP5D*^2^ are mostly expressed in microglia and macrophages. *BIN1* expressed in microglia, oligodendrocyte, and neurons^2^, and *PICALM* expressed in microglia and endothelial cells^2,5^ are associated with endocytosis.

In a normal functional brain, astrocytes, microglia, and oligodendrocytes provide trophic supports to neurons and various cell type specific functions. Astrocytes confer multiple functions to fulfill neurons’ metabolic needs^10^ including but not limited to providing substrates for oxidative phosphorylation^11^, exerting regulation of excitatory CNS neurotransmitter glutamate^12,13^, and serving as bidirectional communication nodes that talk to both neurons and blood vessels and modulate their activities in an arrangement of functional entities named neurovascular units^14^-^16^. Microglia surveil in the extracellular space and look for pathogens or debris to engulf through phagocytosis. Oligodendrocyte provides insulation to neurons by wrapping around the axons with myelin sheath. However, in an AD diseased brain, these supporting cells may become double-edged swords that play beneficial and/or harmful roles as disease progresses. Amyloid-β accumulation and clearance are the central events of the amyloid cascade hypothesis. Both astrocyte and microglia have been involved in response to the toxic stimuli of amyloid plaques. During the early stage, microglia^17-19^ and astrocytes^19-21^ accumulate around plaques to phagocytose or degrade those in a protective manner. However, as disease progresses, the chronic and prolonged activation of microglia and astrocytes will be provoked into a damaging pro-inflammatory state and a vicious circle that exacerbate pathology in a harmful manner. Evidence suggested that increased inflammatory cytokine secretion in microglia, and increased production of complement cascade components, and impaired glutamate regulation (unregulated glutamate activity can cause neuronal excitatory cell death)^13^ may contribute to synaptic loss which ultimately leads to cognitive deficits. Disrupted neuronal plasticity due to myelin loss and dysfunctional neurovascular units further exacerbate the dreadful situation and destroy the harmony of the multi-cell type microenvironment.

Apart from disturbed homeostatic processes and impaired circuits integrity, cell type composition or proportion is also altered. Brains affected by AD exhibits neuronal loss, oligodendrocyte loss, astrocytosis, and microgliosis. However, the specific effects that pathological mutations and risk variants have on brain cellular composition are often ignored. To investigate the changes of cerebral cortex cell-type population structure and account for the associated confounding effects in downstream analysis, we developed an *in-silico* deconvolution method to infer cellular composition from RNA-Seq data, which has been documented in our previous publication^22^. In summary, we firstly assembled a reference panel to model the transcriptomic signature of neurons, astrocytes, oligodendrocytes and microglia. The panel was created by analyzing expression data from purified cell lines. We evaluated various digital deconvolution methods and selected the best performing ones for our primary analyses. We tested the digital deconvolution accuracy on induced pluripotent stem cell (iPSC) derived neurons and microglia, and neurons derived from Translating Ribosome Affinity Purification followed by RNA-Seq. Finally, we verified its accuracy with simulated admixture with pre-defined cellular proportions.

Once the deconvolution approach was optimized, we calculated the cell proportion in AD cases and controls from different brain regions of LOAD and ADAD datasets. We found that neuronal and astrocyte relative proportions differ between healthy and diseased brains, and also differ among AD cases that carry different genetic risk variants. Brain carriers of pathogenic mutations in *APP, PSEN1* or *PSEN2* presented lower neuronal and higher astrocytes relative proportions compared to sporadic AD. Similarly, *APOE* ε4 carriers also showed decreased neuronal and increased astrocyte relative proportions compared to AD non-carriers. In contrast, carriers of variants in *TREM2* risk showed a lower degree of neuronal loss than matched AD cases in multiple independent studies. These findings suggest that different genetic risk factors associated with AD etiology may have gene specific effects in the cellular composition of AD brains.

In a recently published study named PsychENCODE^23^, a very similar deconvolution approach as reported in our previous study^22^ was taken to infer cell type composition from RNA-Seq data predominantly drawn from psychiatric disorder cohorts. From the cell fractions inferred from bulk RNA-Seq data, they found that cell type composition differences can account for more than 88% of bulk tissue expression variation observed across the population with a ±4% variance on a per-subject level. Using cell type compositions as quantitative traits, the authors identified a non-coding variant closed to the *FZD9* gene that is associated with both *FZD9* gene expression and the proportion of excitatory layer 3 neurons^23^. Interestingly, deletion variants found previously upstream of *FZD9* were associated with cell composition changes in Williams syndrome^24^, a developmental disorder exhibits mild to moderate intellectual disabilities with learning deficits and cardiovascular problems. This observation re-emphasized the importance of incorporating cell type composition into RNA-Seq analysis pipeline even in psychiatric disorder cohorts without dramatic changes in cellular composition, not mention the necessity of such practice in neurodegeneration disorders that have significant changes in cell type composition. It also demonstrated the great potential of using relative abundance of specific cell types in identifying novel variants and genes implicated in disease. However, it is unclear if this finding is only applicable to psychiatry-relate traits or it is a more general finding.

In this study, we utilized cell-type proportions inferred from our deconvolution method^22^ to perform cell type QTL analysis in a dataset enriched for AD cases in search for potential new loci that are associated with neurodegeneration disorders. We imputed and merged genotyping or whole genome sequencing data from seven studies - five centered on neurodegeneration (N = 1,125), one schizophrenia cohort (N = 414), and GTEx multiple tissue controls (N = 130). From cortical RNA-Seq data, we derived cell fractions of four major CNS cell types, including neuron, astrocyte, microglia, and oligodendrocyte. Using normalized neuronal proportion as quantitative trait, we identified a variant rs1990621 located in the *TMEM106B* gene region significantly associated with neuronal proportion variation in all cohorts except schizophrenia subjects. This variant is in high LD with rs1990622 (r^2^ = 0.98), which was previously identified as a protective variant in FTD cohorts^25^. Variants in this region have also been found to be associated with AD with TDP-43 pathology^26^, and downregulation of *TMEM106B* is observed in AD brains^27^. In conclusion, we have identified a variant associated with neuronal proportion with potential protective effect in neurodegeneration disorders.

## Methods

### Study participants

The participants were sourced from seven studies with a total sample size of 1,669 (**Table 1**). Among those, five studies are mainly focused on neurodegenerative disorders including Alzheimer’s disease (N = 681), frontotemporal dementia (N = 11), progressive supranuclear palsy (N = 82), pathological aging (N = 29), Parkinson Disease (N = 1), as well as cognitive normal individuals (N = 540). These samples come from the Mayo, MSSM, Knight ADRC, DIAN, and ROSMAP studies as described in **Table 1**. To compare with the neurodegenerative disorders, we also included schizophrenia (N = 210) and bipolar disorders (N = 34) participants from the CommonMind study (**Table 1**). Additionally, two studies, MSSM and GTEx, contain multi-tissue data that include some participants contribute more than one tissue (**Table 1**).

**Table 1.** Demographic information for cohorts included in the study. AD: Alzheimer’s Disease; FTD: frontal temporal dementia; PSP: progressive supranuclear palsy; PA: pathological aging; PD: Parkinson’s Disease; SCZ: schizophrenia; BP: bipolar disease; OTH: other unknown dementia or no diagnosis information.

|  | N | Region | Age | % Male | RIN | TIN | Control | AD | FTD | PSP | PA | PD | SCZ | BP | OTH |
| --- | --- | --- | --- | --- | --- | --- | --- | --- | --- | --- | --- | --- | --- | --- | --- |
| **Discovery** |  |  |  |  |  |  |  |  |  |  |  |  |  |  |  |
| **ROSMAP** | 523 | 1 | 86.6 ± 4.59 | 35.4 | 7.07 ± 0.99 | 73.2 ± 5.13 | 114 | 338 | 0 | 0 | 0 | 0 | 0 | 0 | 71 |
| **Replication** |  |  |  |  |  |  |  |  |  |  |  |  |  |  |  |
| **Mayo** | 260 | 1 | 80.4 ± 8.37 | 48.1 | 8.16 ± 0.903 | 77.4 ± 5.94 | 69 | 80 | 0 | 82 | 29 | 0 | 0 | 0 | 0 |
| **MSSM** | 219 | 4 | 84 ± 7.32 | 35.6 | 6.42 ± 1.77 | 76.4 ± 2.52 | 49 | 170 | 0 | 0 | 0 | 0 | 0 | 0 | 0 |
| **Knight ADRC** | 108 | 1 | 83.1 ± 12 | 42.6 | 6.44 ± 1.2 | 79.4 ± 1.91 | 13 | 77 | 11 | 0 | 0 | 1 | 0 | 0 | 6 |
| **DIAN** | 15 | 1 | 50.9 ± 7.08 | 73.3 | 5.55 ± 1.09 | 78.9 ± 0.99 | 0 | 15 | 0 | 0 | 0 | 0 | 0 | 0 | 0 |
| **GTE<sub>x</sub>** | 130 | 3 | 58.2 ± 9.91 | 67.7 | 6.92 ± 0.846 | 73.8 ± 2.97 | 125 | 1 | 0 | 0 | 0 | 0 | 0 | 0 | 4 |
| **CommonMind** | 414 | 1 | 64.6 ± 18 | 62.3 | 7.67 ± 0.899 | 50 ± 7.21 | 170 | 0 | 0 | 0 | 0 | 0 | 210 | 34 | 0 |
| **Replication Total** | 1,146 |  |  |  |  |  | 426 | 343 | 11 | 82 | 29 | 1 | 210 | 34 | 10 |
| **Merged Total** | 1,669 |  |  |  |  |  | 540 | 681 | 11 | 82 | 29 | 1 | 210 | 34 | 81 |

### Standard protocol approvals, registrations and patient consents

The protocol of DIAN and Knight-ADRC studies have been approved by the review board of Washington University in St. Louis. The protocol of Mayo dataset was approved by the Mayo Clinic Institutional Review Board (IRB). All neuropsychological, diagnostic and autopsy protocols of MSSM dataset were approved by the Mount Sinai and JJ Peters VA Medical Center Institutional Review Boards. The religious orders study and the memory and aging project of ROSMAP was approved by the IRB of Rush University Medical Center. The NIH Common Fund’s GTEx program protocol was reviewed by Chesapeake Research Review Inc., Roswell Park Cancer Institute’s Office of Research Subject Protection, and the institutional review board of the University of Pennsylvania. Within CommonMind consortium, the MSSM sample protocol was approved by Icahn School of Medicine at Mount Sinai IRB; the Pitt sample protocol was approved by the University of Pittsburgh’s Committee for the Oversight of Research involving the Dead and IRB for Biomedical Research; the Penn sample protocol was approved by the Committee on Studies Involving Human Beings of the University of Pennsylvania. All participants were recruited with informed consent for research use.

### Data collection and generation

Cortical tissues from various locations of post-mortal brains were collected (**Table 2**). RNA was extracted from lysed tissues and prepared into libraries of template molecules ready for subsequent next-generation sequencing steps. Ribosomal RNAs constitute 80%-90% of total RNAs, which are not the targets of this study. To focus on mRNA quantification usually researchers would either remove excessive rRNAs or enrich for mRNAs during RNA-Seq library preparation. In this study, DIAN^22^, Knight ADRC^22^, MSSM^28^, and CommonMind^29^ took a rRNA depletion approach to removed ribosomal RNA from total RNAs to retain a higher mRNA content. Whereas, Mayo^30^, ROSMAP^31,32^, and GTEx^33,34^ took a poly-A enrichment approach to enrich mRNAs from total RNAs. Genotype information were also collected and sequenced correspondingly. RNAseq paired with genotype data for each participant were either sequenced at Washington University for DIAN and Knight-ADRC studies or downloaded from public database for all the other studies. Please see **Table 2** and each study reference(s) for more data collection and generation specifications.

**Table 2.** General information of seven studies evolved in the analysis. TCX: temporal cortex; PAR: parietal cortex; CTX: cortex; FCX: frontal cortex; DLPFC: dorsal lateral prefrontal cortex. BM9: dorsal lateral prefrontal cortex; BM10: Anterior prefrontal cortex; BM22: superior temporal gyrus; BM24: ventral anterior cingulate cortex; BM36: parahippocampal gyrus; BM44: inferior frontal gyrus. Mean coverage unit is million.

| Dataset | Brain Region | Library Type | Read Length | mRNA Enrichment | Sequencer | Mean Coverage | DNA type | Reference |
| --- | --- | --- | --- | --- | --- | --- | --- | --- |
| **Discovery** |  |  |  |  |  |  |  |  |
| **ROSMAP** | DLPFC | Paired end | 101 | poly-A selection | HiSeq 2000 | 99.2 ± 29.29 | WGS | Bennett 2012; Bennett 2012 |
| **Replication** |  |  |  |  |  |  |  |  |
| **Mayo** | TCX | Paired end | 101 | poly-A selection | HiSeq 2000 | 158.31 ± 34.04 | Genotype | Allen 2016 |
| **MSSM** | BM10 BM22<br>BM36 BM44 | Single end | 100 | rRNA depletion | HiSeq 2500 | 35.96 ± 10.04 | WGS | Wang 2018 |
| **Knight-ADRC** | PAR | Paired end | 150 | rRNA depletion | HiSeq 4000 | 137.87 ± 21.81 | Genotype | Li 2018 |
| **DIAN** | PAR | Paired end | 150 | rRNA depletion | HiSeq 4000 | 149.82 ± 19.68 | Genotype | Li 2018 |
| **GTE<sub>x</sub>** | BM24 CTX<br>FCX | Paired end | 76 | poly-A selection | HiSeq 2000 | 48.28 ± 13.2 | WGS | GTE<sub>x</sub> 2013; Battle 2017 |
| **Common Mind** | BM9 | Paired end | 100 | rRNA depletion | HiSeq 2500 | 86 ± 21.12 | Genotype | Fromer 2016 |

### Data QC and preprocessing

#### Genetic Data

Stringent quality control (QC) steps were applied to each genotyping array or sequence data. The minimum call rate for single nucleotide polymorphisms (SNPs) and individuals was 98% and autosomal SNPs not in Hardy-Weinberg equilibrium (p-value < 1×10^−06^) were excluded. X-chromosome SNPs were analyzed to verify gender identification. Unanticipated duplicates and cryptic relatedness (Pihat ≥ 0.25) among samples were tested by pairwise genome-wide estimates of proportion identity-by-descent. EIGENSTRAT^35^ was used to calculate principal components. The 1000 Genomes Project Phase 3 data (October 2014), SHAPEIT v2.r837^36^, and IMPUTE2 v2.3.2^37^ were used for phasing and imputation. Individual genotypes imputed with probability < 0.90 were set to missing and imputed genotypes with probability ≥0.90 were analyzed as fully observed. Genotyped and imputed variants with MAF < 0.02 or IMPUTE2 information score < 0.30 were excluded. WGS data quality is censored by filtering out reads with sequencing depth DP < 6 and quality GQ < 20 followed by similar QC approaches as described above for genotyping data. After the QC, all studies including imputed genotype and WGS data was merged into a binary file using Plink for downstream analysis. PCA and IBD analyses were performed on the merged binary files using Plink to keep European ancestry and unrelated participants (**Figure 1** and **Figure 2**).

**Figure 1.**
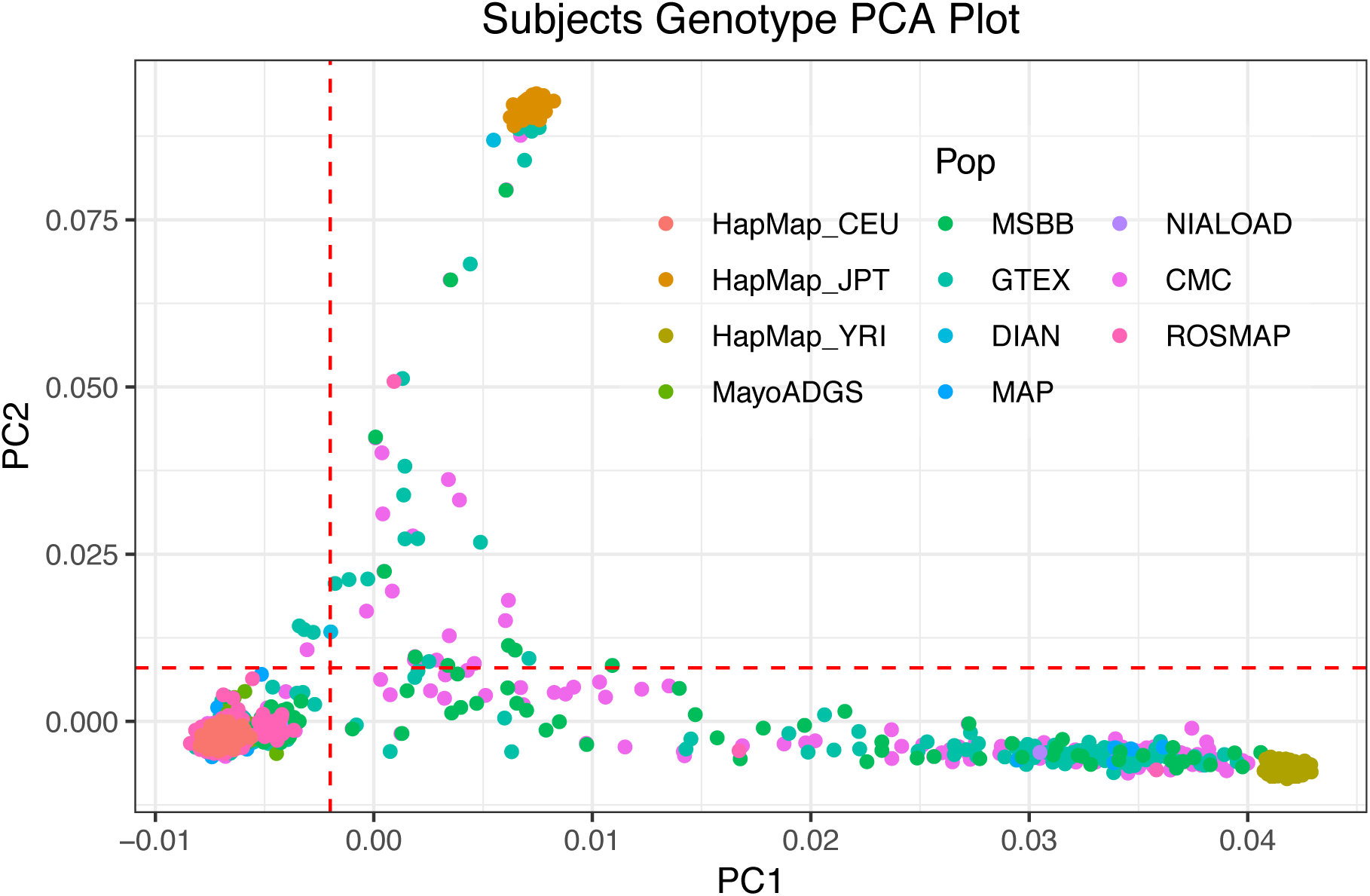
Genomic PCA analysis. Genotype data PCA analysis was performed to select European ancestry subjects with PC1 < −0.002 and PC2 < 0.008 with red dotted cut-off lines. HapMap_CEU: HapMap Utah residents with Northern and Western European ancestry; HapMap_JPT: HapMap Japanese in Tokyo, Japan; HapMap_YRI: HapMap Yoruba in Ibadan, Nigeria; MayoADGS: Mayo Clinic study participants; MSBB: MSSM study participants; GTEX: GTEx study participants; DIAN: DIAN study participants; MAP: Knight-ADRC participants; NIALOAD: Knight-ADRC participants; CMC: CommonMind participants; ROSMAP: ROSMAP participants.

**Figure 2.**
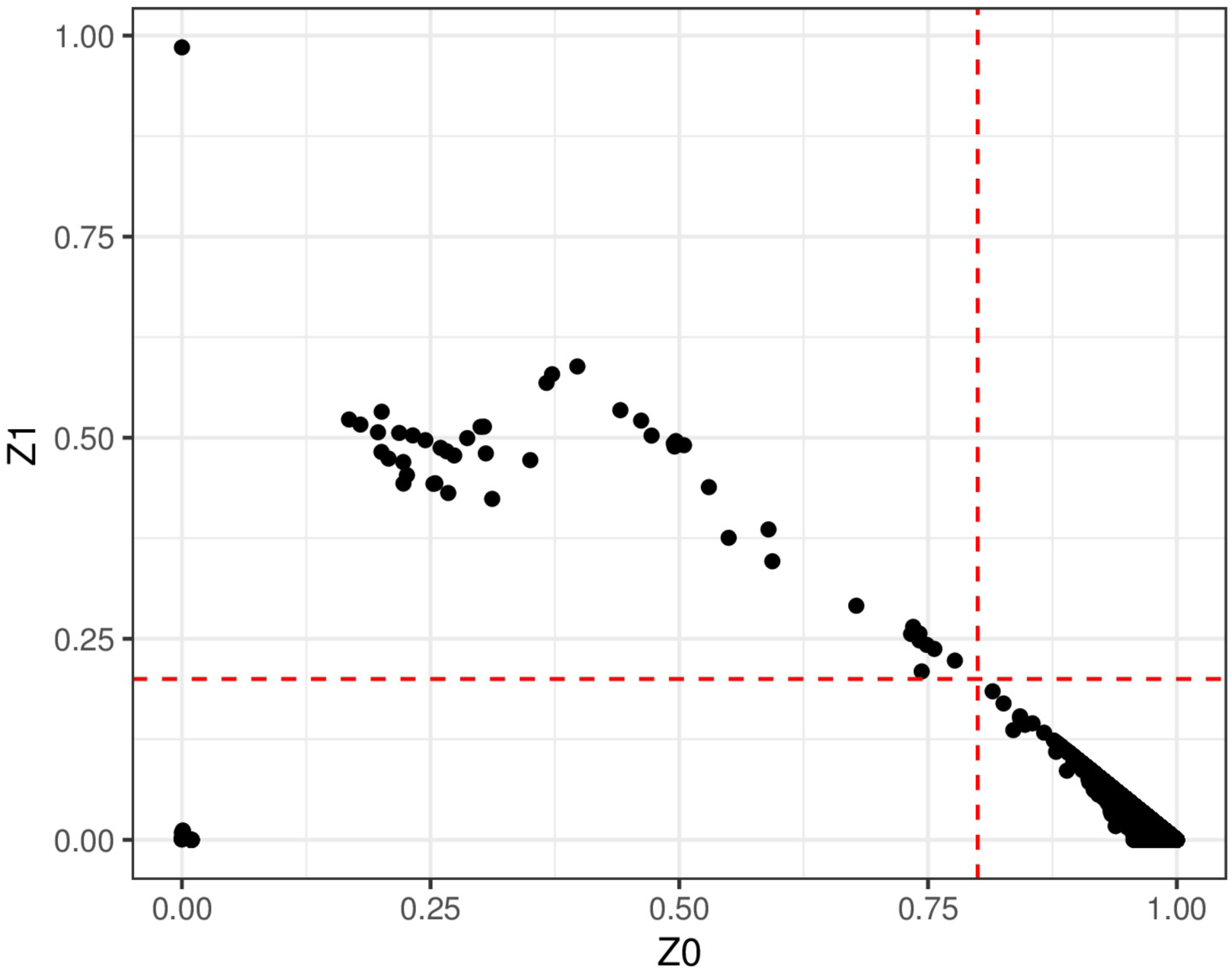
Genomic IBD analysis. IBD analysis was performed to select unrelated subjects with Z0 > 0.8 and Z1 < 0.2 with red dotted cut-off lines. When there are related individuals, one individual will be dropped from the related pair.

#### Expression Data

FastQC was applied to RNAseq data to examine various aspects of sequencing quality^38^. Outlier samples with high rRNA contents or low sequencing depth were removed from the pool. The remaining samples were aligned to human GRCh37 primary assembly using Star with 2-Pass Basic mode (ver 2.5.4b)^39^. Alignment metrics were ascertained by applying Picard CollectRnaSeqMetrics^40^ including reads bias, coverage, ribosomal contents, coding bases, and etc. Following which, transcript integrity number (TIN) for each transcript was calculated on aligned bam files using RSeQC tin.py^41^ (ver 2.6.5). RNAseq coding gene and transcript expression was quantified using Salmon transcript expression quantification (ver 0.7.2) with GENCODE *Homo sapiens* GRCh37.75 reference genome^42^.

Four major central nerve system cell type proportions were inferred from RNAseq gene expression quantification output as documented in our previous deconvolution study^22^. To briefly explain the deconvolution process, we firstly assembled a reference panel to model the transcriptomic signature of neurons, astrocytes, oligodendrocytes and microglia from purified single cell tissue sources respectively. Using the reference panel and the method population-specific expression analysis^43^ (PSEA, also named meanProfile in CellMix implementation^44^), we calculated four cell type proportions for each subject bulk RNAseq data. For each brain tissue collection site of each study, outlier values for each cell type proportion were removed. Mean values for each cell type of each tissue in each study were subtracted from the deconvolution results to center all the distributions to zero mean (**Figure 3**). Phenotype information from all studies were merged and unified to the same coding paradigm to enable downstream joint analysis; for example, males are all coded as 1 and females are 2.

**Figure 3.**
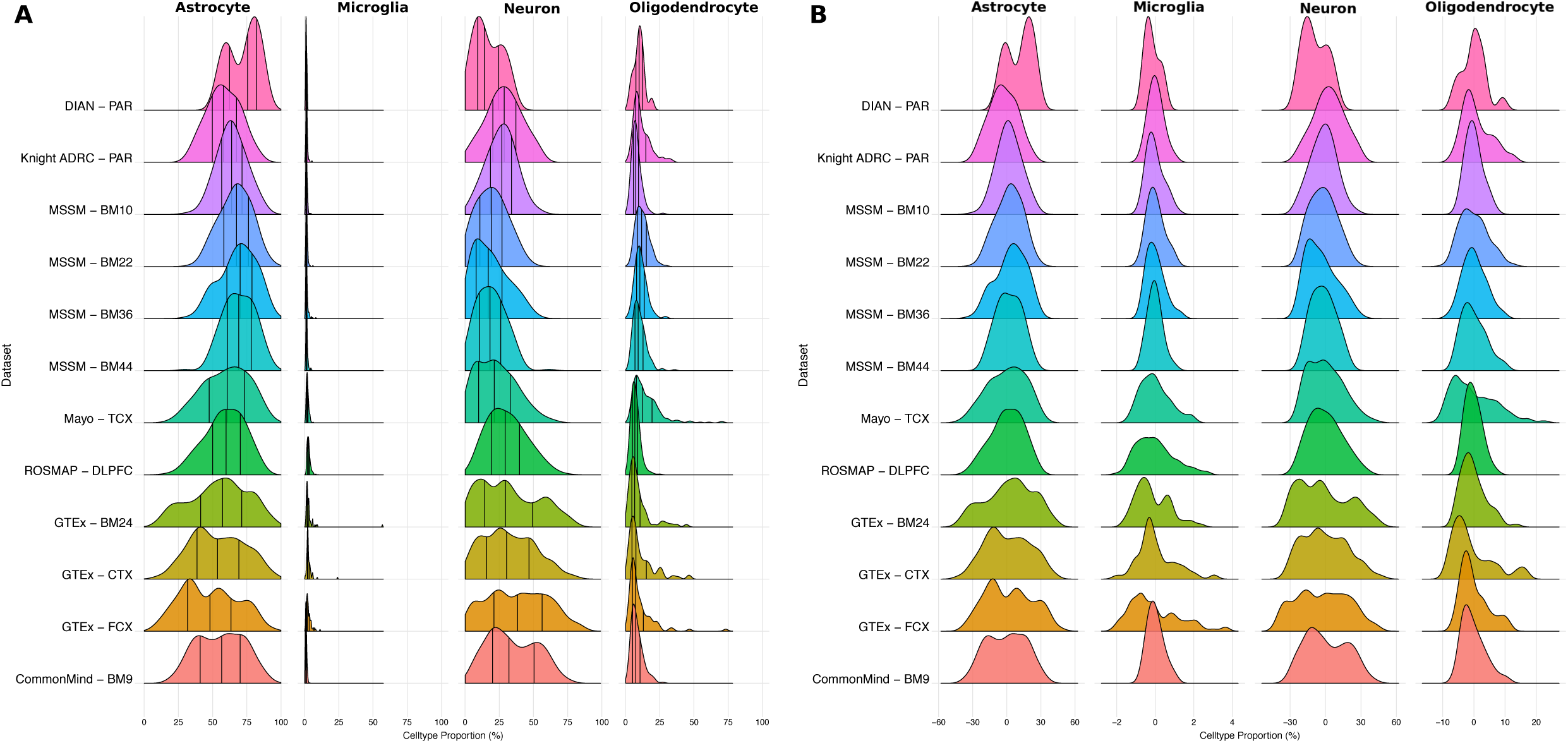
Cell proportion distribution. Major CNS cell type proportions derived from RNAseq datasets with each row representing each tissue of each study A) raw cell type proportions inferred from the data with vertical bars indicating quantiles within each tissue and each cell type. B) cell type proportions were normalized by subtracting the mean from each tissue deconvolution result after removing outliers.

### Data analysis

For the discovery phase, ROSMAP dataset was analyzed with linear regression model employed in Plink^45^ using normalized neuronal proportion to run quantitative trait analysis. Age, sex, PC1, PC2, and median TIN were used as covariates to account for potential genetic, phenotypic or technical heterogeneity. TIN is calculated directly from post-sequencing results that captures RNA degradation by measuring mRNA integrity directly^41^. Results were depicted as Manhattan plots using R (ver 3.4.3) qqman package^46^ (ver 0.1.4).

For the replication phase, all the other studies except ROSMAP were combined and prepressed to run meta-tissue QTL analysis because MSSM and GTEx contain samples with multiple cortical tissues. Meta-Tissue software installation and data preprocessing were conducted following a four-step instruction documented in the developer website: http://genetics.cs.ucla.edu/metatissue/install.html. Meta-tissue^47^ processing pipeline calls two main functions, firstly MetaTissueMM^47^ and then followed by Metasoft^48^. MetaTissueMM applies a mixed model to account for the heterogeneity of multiple tissue QTL effects. Metasoft performs the meta-analysis while proving a more accurate random effect p-value for multiple tissue analysis and a m-value based on Bayesian inference to indicate how likely a locus is a QTL in each tissue. Similarly, results were depicted as Manhattan plots and visually examined.

For the final merging phase, both discovery and replication studies were combined to maximize sample size. Apart from meta-tissue analysis by each tissue of each study, a split by disease status analysis was also performed in the final merging phase. Samples from each tissue of each study were also split into disease categories. Resultant subcategory with less than 20 subjects were removed from the analysis to avoid false results due to too small sample size. Similar data preparation and analysis pipeline were enforced as documented above.

QTL analysis results were uploaded to Fuma (v1.3.3d)^49^ to annotation significant SNPs (p-value < 10^−06^) with GWAScatalog (e91_r2018-02-06) and ANNOVAR (updated 2017-07-17). Gene-based analysis was also performed by Magma (v1.06)^50^ implemented in Fuma.

### Data availability

Mayo: https://www.synapse.org/#!Synapse:syn5550404

MSSM: https://www.synapse.org/#!Synapse:syn3157743

ROSMAP: https://www.synapse.org/#!Synapse:syn3219045

CommonMind: https://www.synapse.org/#!Synapse:syn2759792

GTEx: https://www.ncbi.nlm.nih.gov/projects/gap/cgi-bin/study.cgi?study_id=phs000424.v7.p2

Knight-ADRC: https://www.synapse.org/#!Synapse:syn12181323

According to the data request terms, DIAN data are available upon request: http://dian.wustl.edu

## Results

### tudy Design

The ROSMAP study containing 523 subjects will be the discovery dataset, and the other six studies are collapsed into replication dataset with 1,146 subjects. Altogether, we have assembled a set of cortical RNA-Seq data comprised of 1,669 participants predominantly focused on neurodegenerative disorders from seven sources (**Figure 4**, **Table 1**). Collectively, Mayo, MSSM, Knight ADRC, and ROSMAP studies contributed 664 sporadic AD cases. Apart from sporadic AD, 15 subjects from DIAN study and 2 from Knight-ADRC also harbor *PSEN1, PSEN2*, and *APP* mutations that exhibit familial AD inheritance pattern. Other neurodegenerative disorders, including progressive supranuclear palsy (PSP), pathological aging (PA), frontal temporal dementia (FTD), and Parkinson’s Disease (PD), are mainly drawn from Mayo and Knight ADRC datasets. Other psychiatric disorders including schizophrenia and bipolar disorders are contributed by the CommonMind study. Besides, 540 control subjects cleared of cognitive dementia or neuropsychiatric symptoms were also included. MSSM and GTEx also included multiple tissue data, which were collected from multiple regions of the same subjects that allow us to perform region specific comparison within the same cohort.

**Figure 4.**
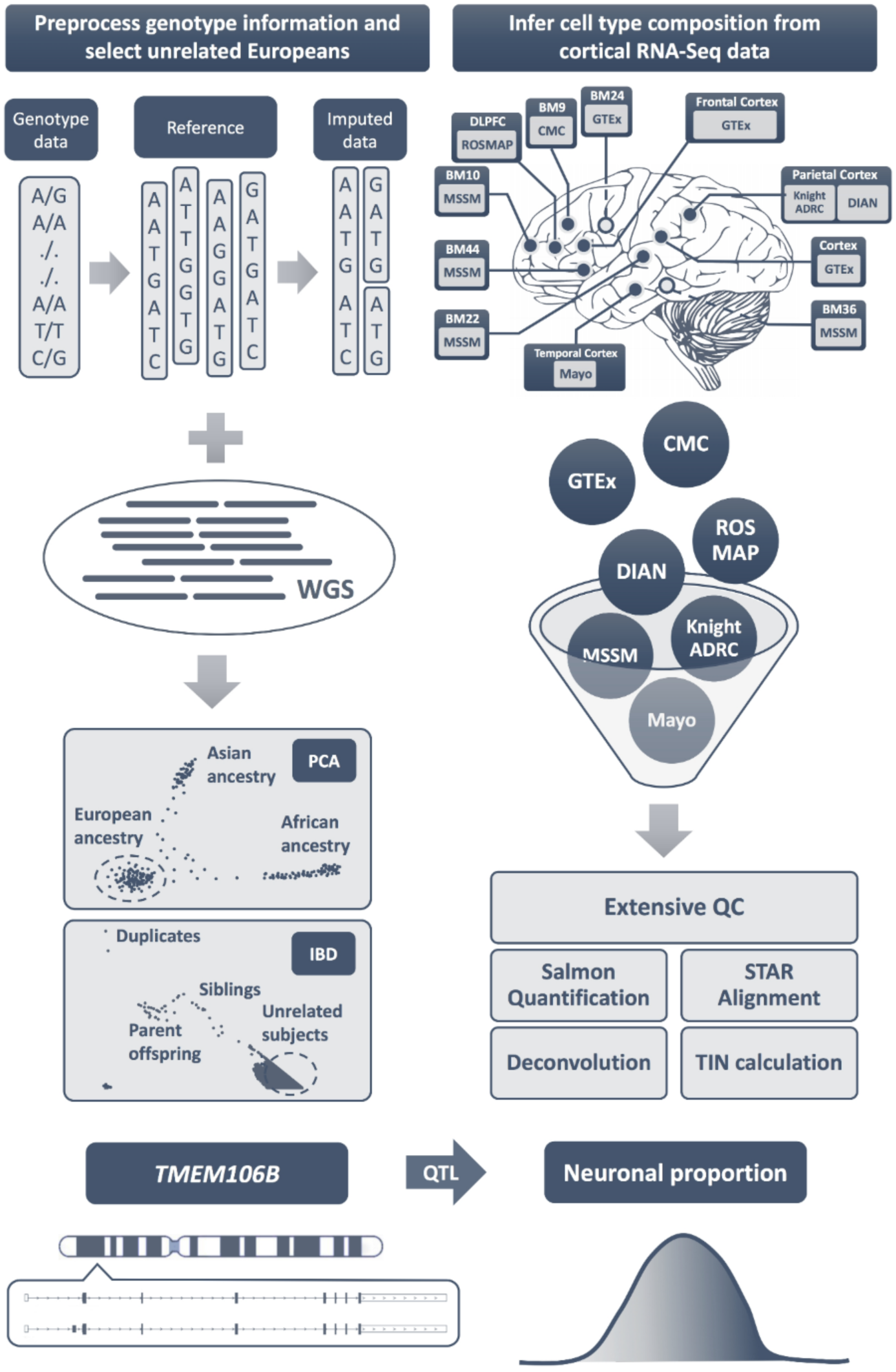
Study Design. RNA-Seq and paired genotype or WGS data were accessed and preprocessed for downstream analysis. Genotype data was censored based on our quality control criteria and imputed as needed. WGS and imputed genotype were merged and followed by PCA and IBD procedures to select unrelated European ancestry subjects. RNA-Seq data was quality checked with FastQC and aligned to human GRCh37 primary assembly with Star, from which TIN was inferred with RSeQC to account for RNA integrity variances that we later incorporated into the analysis. Gene expression were quantified from unaligned RNA-Seq with psedo-aligner Salmon for deconvolution procedure. Cell type composition comprised of four major CNS cell type proportions were inferred by performing deconvolution procedure on gene expression quantification results. Using cell type proportions as quantitative traits, we identified loci in *TMEM106B* gene region associated with neuronal proportion in our assembled dataset.

Discovery analysis was performed in ROSMAP study. In the replication phase, all the other studies were merged to replicate signals identified from the discovery ROSMAP set. Because GTEx and MSSM contain multiple cortical regions collected from the same subjects, we also applied meta-tissue software^47^ specifically designed for multi-tissue QTL analysis to perform a mixed model analysis with random effects that account for correlated measurements from multi-tissue individuals. To attain the largest available sample size for this study, the discovery and replication sets were merged to perform the merged multi-tissue QTL analysis in a search for additional signals hidden in previously separated discovery or replication analysis due to lack of power. After merged analysis, the cohorts were split into four major disease status groups (AD, control, schizophrenia, other non-AD neurodegenerative disorders) to explore how different disease strata could impact the results.

### TMEM106B variants associated with neuronal proportion

During discovery phase, ROSMAP dataset (N = 484 after removing outliers from total number of 523 subjects) was used to perform cell type proportion QTL analysis. Using normalized neuronal proportion as a quantitative trait, the QTL analysis identified more than 10 peaks that passed genome wide suggestive threshold (<1.0×10^−06^, **Figure 5AB, Table 3**). However, only one signal rs1990621 (chr7: 12283873) were replicated with p-value = 7.41×10^−04^ in the replication dataset (N = 1,052) combining all the other datasets except ROSMAP (**Figure 5CD**). When the discovery and replication datasets were merged to attained a larger sample size (N = 1,536), rs1990621 major allele C is negatively associated with neuronal proportion with p-value = 9.42×10^−09^ (**Figure 6AB, Figure 7AC**), which means the minor allele G is associated with increased neuronal proportion in our assembled datasets focusing on neurodegenerative disorders.

**Table 3.** rs1990621 (chr7:12283873) major allele C is significantly associated with decreased neuronal proportions. Therefore, G allele (MAF = 0.4658) is significantly associated with increased neuronal proportions.

| Dataset | Brain Region | Ref Allele | Sample Size | Beta | SE | P-value |
| --- | --- | --- | --- | --- | --- | --- |
| Discovery | DLPFC | C | 484 | -0.3 | 0.06 | $6.40 \times 10^{-07}$ |
| Replication | Multiple | C | 1,052 | -0.13 | 0.04 | $7.41 \times 10^{-04}$ |
| Merged meta-tissue | Multiple | C | 1,536 | -0.16 | 0.05 | $9.42 \times 10^{-09}$ |

**Figure 5.**
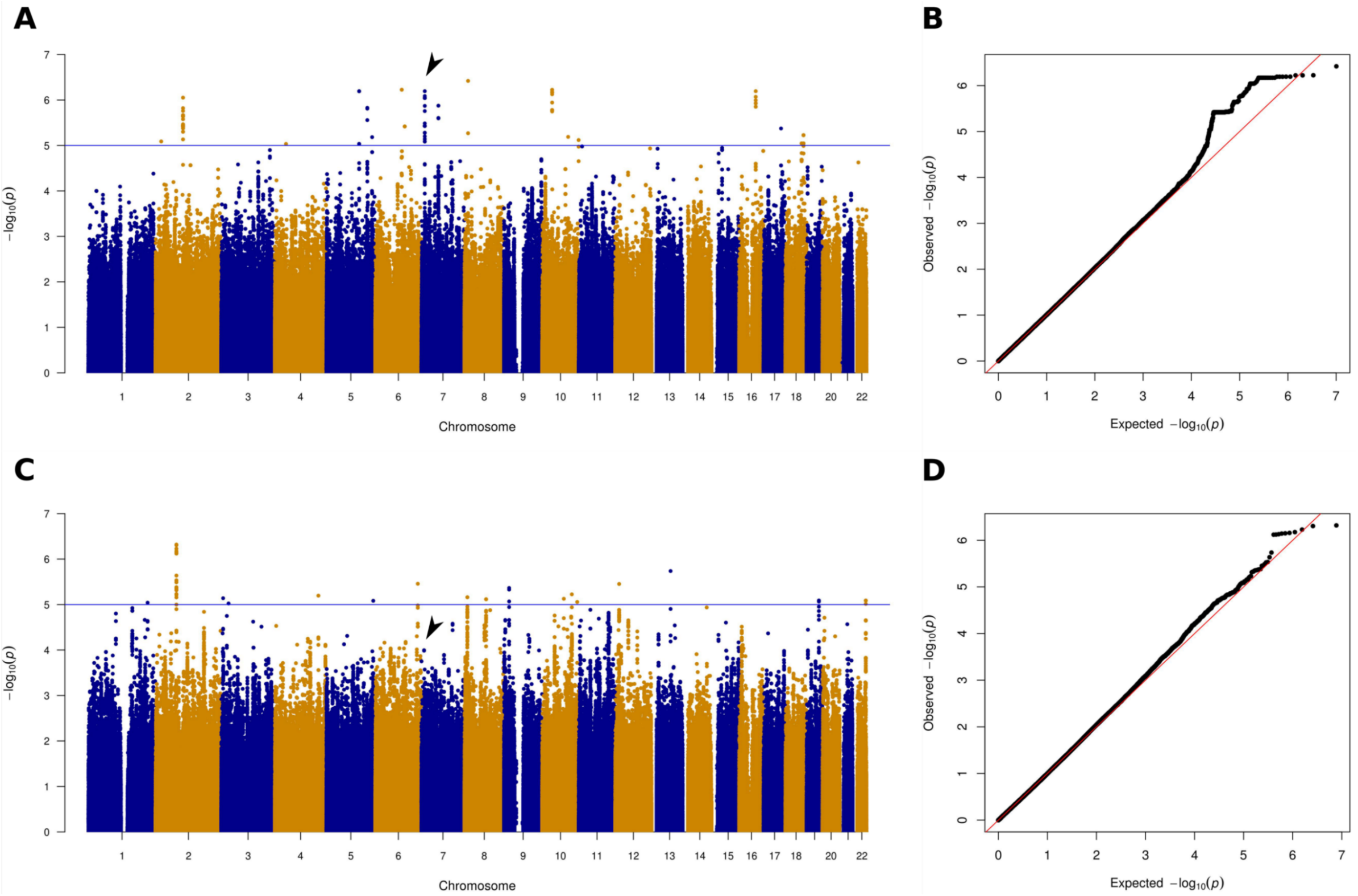
Discovery and replication phases Manhattan and QQ plots. Loci located in chromosome 7 were associated with neuronal proportion in ROSMAP discovery dataset and replicated in replication dataset. A) Discovery set Manhattan plot showed seven peaks associated with neuronal proportion at suggestive threshold. The peak located in chromosome 7 was labeled, which is for rs1990621 with p-value = 6.4×10^−07^. B) QQ plot of the discovery phase analysis. C) Replication set Manhattan plot showed that the peak located in chromosome 7 replicated the signal identified during discovery phase with p-value = 7.41×10^−04^. D) QQ plot of the replication phase analysis.

**Figure 6.**
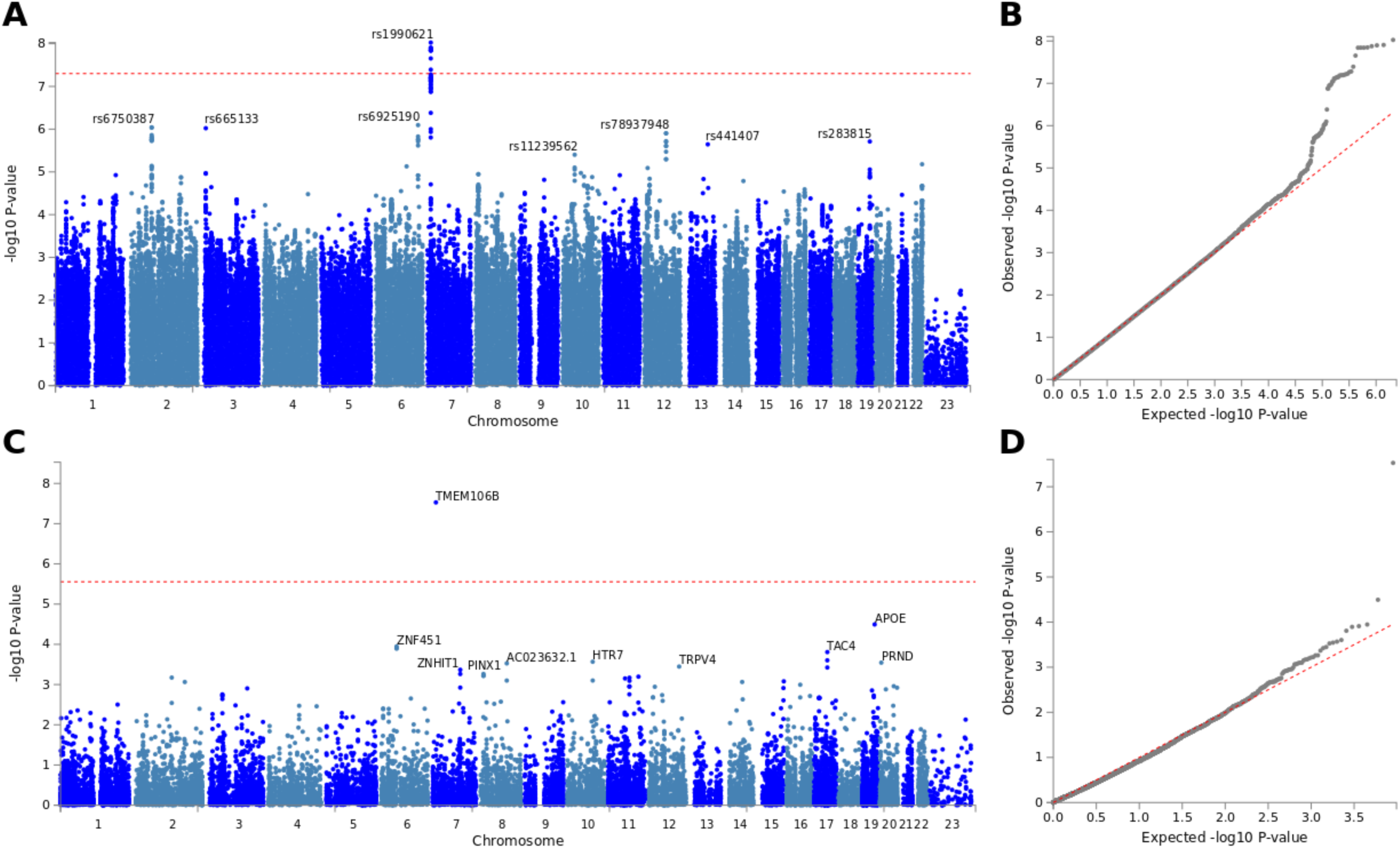
Merged SNP-based and gene-based analysis. rs1990621 located in chromosome 7 *TMEM106B* gene region was significantly associated with neuronal proportion in cortical RNAseq dataset. A) Manhattan plot showed SNP-based genome-wide significant hit located in chromosome 7 with other suggestive SNP hits labeled. B) QQ plot of the SNP-based analysis. C) Manhattan plot showed gene-based genome-wide significant hit located in chromosome 7 with other suggestive gene hits labeled. D) QQ plot of the gene-based analysis.

**Figure 7.**
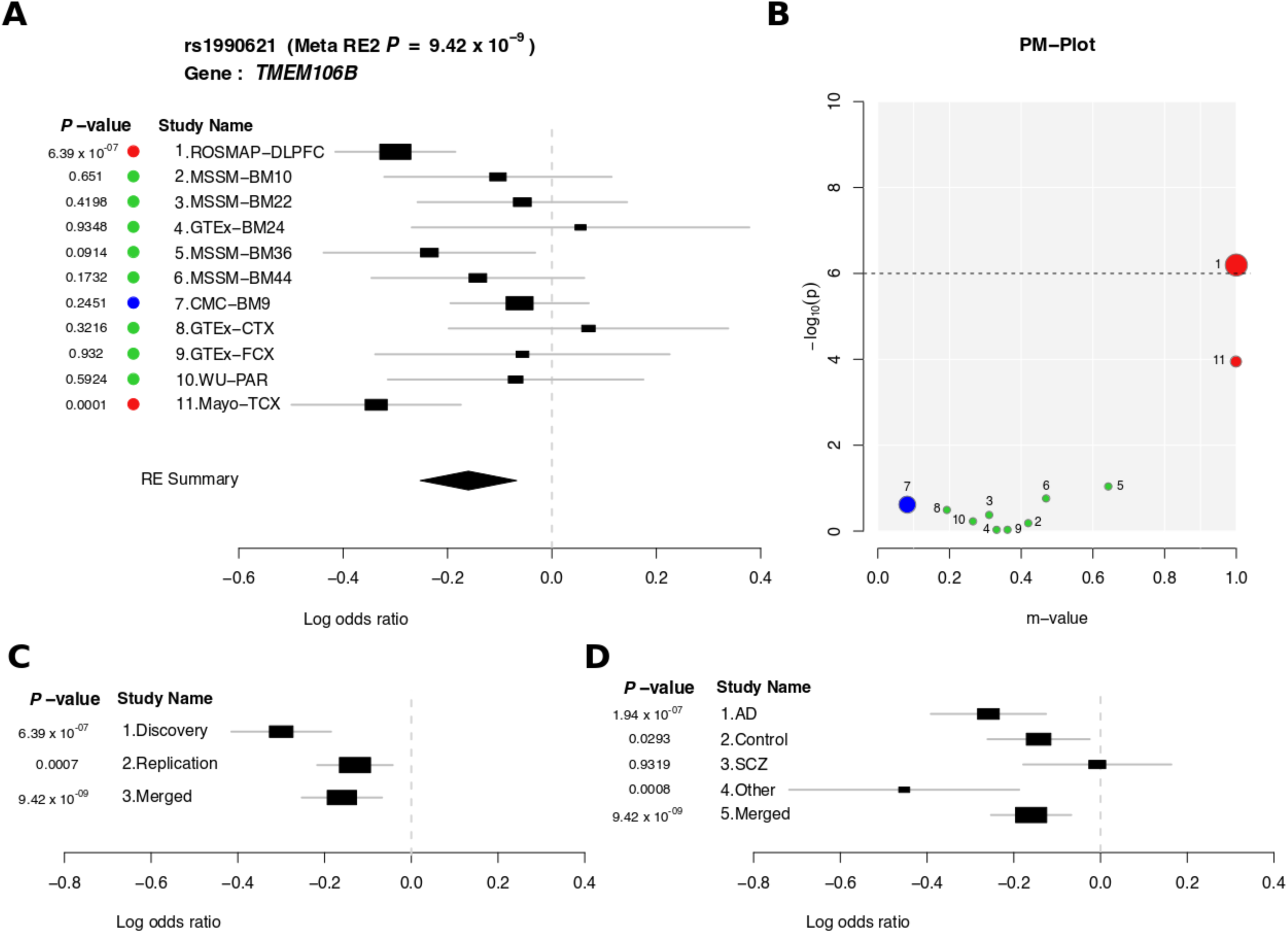
Meta-Tissue analysis results of rs1990621. A) Forest plot showed p-value and confidence interval for rs1990621 for each tissue site of each dataset that included in the Meta-Tissue analysis. Summary random effect was depicted at the bottom as RE Summary. B) PM-Plot of rs1990621 while combining both p-value (y axis) and m-value (x axis). Red dot indicates that the variant is predicted to have an effect in that particular dataset, blue dot means that the variant is predicted to not have an effect, and green dot represents ambiguous prediction. C) Forest plot p-value and confidence interval for rs1990621 for discovery, replication, and merged datasets. D) Forest plot p-value and confidence interval for rs1990621 when splitting the merged dataset into four main disease categories.

Noticeably, in both replication and merged analyses, multi-tissue data were involved that provided additional power but also posed challenges to the analysis, the same issue faced by the GTEx study^34,51^. Compared to a tissue-by-tissue approach, multiple tissues collected from the same subject may help identify QTL by aggregating evidence from multiple tissues, which is similar to a meta-analysis of combining each study. However, one violation of such approach is that the tissues collected from the same subject are presumably highly correlated since they shared the same genetic architecture. Thus, it violates the assumption of independency for carrying out a standard meta-analysis^47^. Another challenge of the multi-tissue QTL is the heterogeneity of the effects, which means a variant may have different effects on different tissues. To resolve these issues, we applied the Meta-Tissue analytic pipeline^47^ (http://genetics.cs.ucla.edu/metatissue/) specifically designed for multi-tissue QTL, the same approach that GTEx took to analyze their multi-tissue data. As shown in **Figure 7A**, Meta-Tissue analysis results of rs1990621 for the merged analysis were displayed as a forest plot with 95% confidence interval and p-value labeled for each tissue of each study. Among them, MSSM and GTEx are multi-tissue studies while the others are single-tissue studies. Meta-Tissue used a linear mixed model to capture the multi-tissue correlation within MSSM and GTEx respectively. Regarding the effect heterogeneity, Meta-Tissue calculated a m-value^48^ to predict if a variant has an effect in a tissue. M-value is similar to the posterior probability of association based on the Bayes factor^48^ but with differences specifically designed for detecting whether an effect is present in a study included in a meta-analysis. **Figure 7B** is a PM-Plot that integrates evidences from both frequentist (p-value) and Bayesian (m-value) sides to interpret the heterogeneity of multi-tissue QTL effects. Variant rs1990621 in ROSMAP and Mayo studies have m-values greater than 0.9, are predicted to have an effect and color coded with red. In CMC study, the m-value is less than 0.1, so it is predicted to not to have an effect and color coded with blue. All the other studies with m-value between 0.1 and 0.9 are predicted with ambiguous effect and color coded with green. Based on the forest plot and PM-Plot, the variant does have effect heterogeneity across different tissues and studies. In this case, random-effect model will be more suitable to account for effect heterogeneity. Therefore, summary random effect and p-value were reported for the analysis.

Apart from multi-tissue QTL, a single-tissue joint analysis was also performed. In this case, one tissue region was drawn from the multi-tissue data to avoid violating the independency assumption. Specifically, BM36 and frontal cortex tissue were selected to represent MSSM and GTEx study respectively. Study sites were coded as dummy variables to account for potential batch effects. In this joint analysis, the variant rs1990621 is also the top hit with p-value = 7.66×10^−10^.

### Neuronal protective effect of TMEM106B variants observed in neurodegenerative disorders and normal aging participants

To explore the effect in different disease categories, the merged dataset was stratified based on disease: AD, other non-AD neurodegenerative disorders, schizophrenia and control. Signification associations between rs1990621 and neuronal proportion were observed in AD (p-value = 1.95×10^−07^), other non-AD neurodegenerative (p-value = 8.19×10^−04^), and cognitive normal control (p-value = 2.94×10^−02^) cohorts, but not in schizophrenic cohort (p-value = 9.32×10^−01^, **Table 4**, **Figure 7D**). The effect of the variant was more prominent in neurodegenerative cohorts and aging controls with mean age of death greater than 65 years old. However, it was absent from younger cohorts such as GTEx controls and CommonMind schizophrenia participants. Thus, this variant seems to be associated with a neuronal protection mechanism shared by any aging process in the present or absence of neuropathology.

**Table 4.** rs1990621 (chr7:12283873) major allele C is significantly associated with decreased neuronal proportions in AD, Control, and other non-AD neurodegenerative disorders. SCZ: schizophrenia; other: other non-AD neurodegenerative disorders, including progressive supranuclear palsy and pathological aging. BM9: dorsal lateral prefrontal cortex. TCX: temporal cortex.

| Disease | Brain Region | Ref Allele | Sample Size | Beta | SE | P-value |
| --- | --- | --- | --- | --- | --- | --- |
| AD | Multiple | C | 639 | -0.26 | 0.07 | $1.95 \times 10^{-07}$ |
| Control | Multiple | C | 476 | -0.14 | 0.06 | $2.94 \times 10^{-02}$ |
| SCZ | BM9 | C | 189 | -0.01 | 0.09 | $9.32 \times 10^{-01}$ |
| Other | TCX | C | 103 | -0.45 | 0.14 | $8.19 \times 10^{-04}$ |

### Functional annotation

The variant rs1990621 is located in the *TMEM106B* gene region where other variants in high LD linkage are also located and labeled in **Figure 8A**. Although the CADD score and RegulomeDB score for this variant are not remarkably high to suggest any functional consequences (**Figure 8BC**), this variant is in high LD with rs1990622 (r^2^ = 0.98), a *TMEM106B* variant previous identified to be associated with FTD risk^25^, particularly in granulin precursor (GRN) mutation carriers^52,53^. *TMEM106B* is expressed in neurons and microglia, with highest protein expression detected in the late endosome/lysosome compartments of neurons^54-57^. A nonsynonymous variant rs3173615, which is also in high LD with rs1990621 (r^2^ = 0.98), located in the exon 6 of *TMEM106B* (the dark blue dot in **Figure 8B**) produces two protein isoforms (p.T185S) that affect TMEM106B protein level through protein degradation mechanism^55,58,59^.

**Figure 8.**
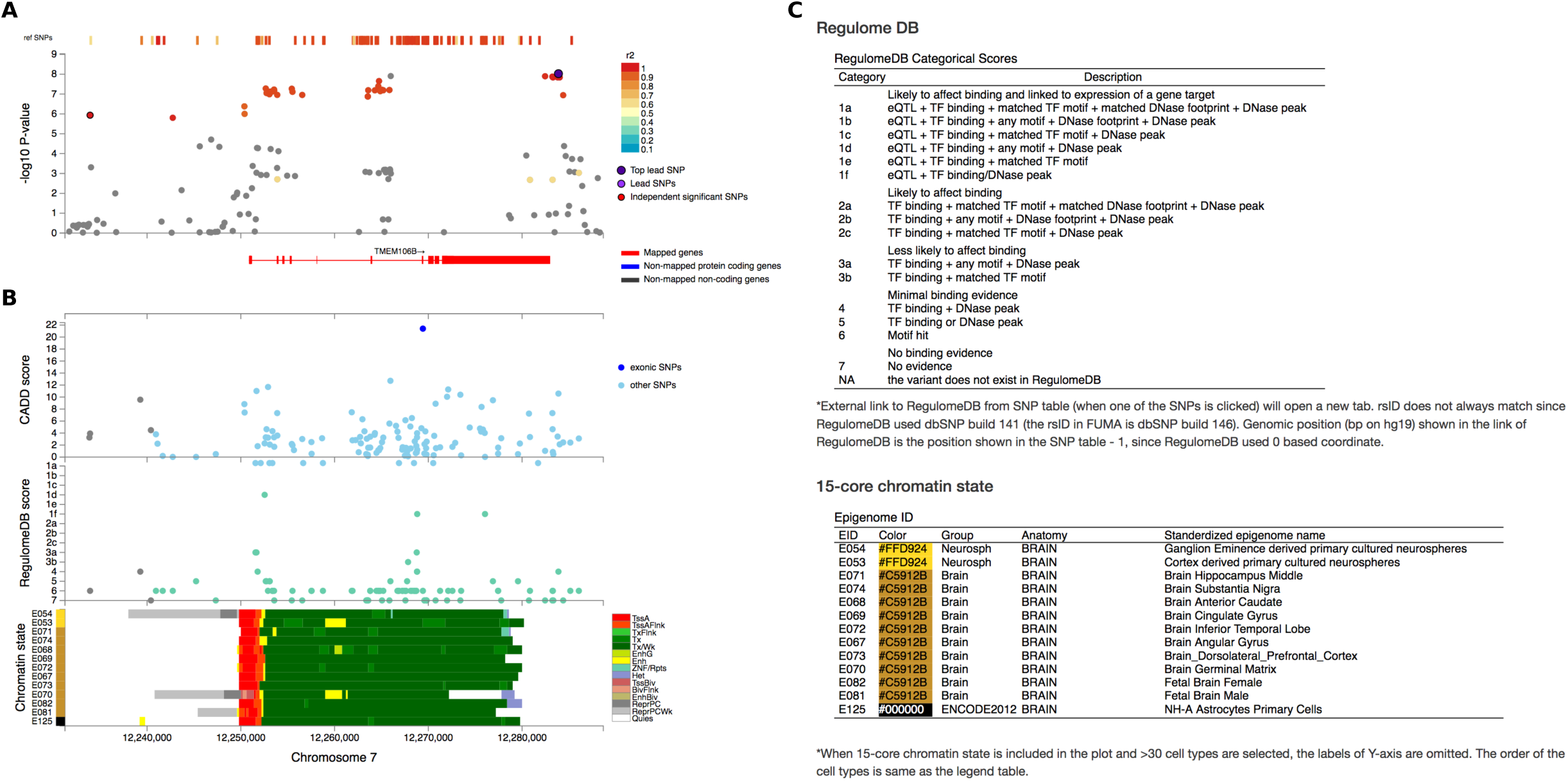
Variant rs1990621 functional annotation and local plot. A) Local plot showed the zoom-in view of the hit in chromosome 7 with the top lead SNP rs1990621 labeled with dark purple. Nearby SNPs were also mainly located in the *TMEM106B* gene region and color coded with LD r2 thresholds. B) Bottom panel showed combined CADD score, RegulomeDB score, and Chromatin state of the region shown in the top panel. C) Regulome DB and chromatin state explanation.

### The impact of other neurodegenerative risk loci on neuronal proportion

To investigate what other AD or FTD variants might have an effect in neuronal proportion QTL analysis, we extracted results for 38 SNPs examined in two large scale genome wide association studies, AD focused (Lambert et al.^2^) and FTD focused (Ferrari et al.^60^) studies. Among those, only variants located in *TMEM106B* and *APOE* gene regions passed genome wide significant or suggestive threshold. Both rs1990622 (**Figure 9A**) and rs2075650 (**Figure 9B**) were found to be associated with FTD reported in Ferrari et al., which were associated with neuronal proportion in this study (**Table 5**). The top signals in *APOE* region are rs283815, rs769449, and rs429358 with p-value < 1.22×10^−05^. Note that rs429358 is one of the two SNPs that determine *APOE* isoforms. Remember that *APOE* ε4 alleles, coded by rs429358(C) and rs7412(C), confers the largest effect for AD risk. We observed that the C allele of rs429358 was associated with decreased neuronal proportion, but no association observed between rs7412 and neuronal proportion.

**Table 5.** Neuronal proportion cQTL p-values were reported for variants previously identified in AD risk (by Lambert et al.) and FTD risk (by Ferrari et al.) studies.

| SNP | CHR | BP | cQTL p-value | AD risk p-value | FTD risk p-value |
| --- | --- | --- | --- | --- | --- |
| rs6656401 | 1 | 207,692,049 | $5.59 \times 10^{-02}$ | $5.7 \times 10^{-24}$ | - |
| rs35349669 | 2 | 234,068,476 | $9.80 \times 10^{-01}$ | $3.2 \times 10^{-08}$ | - |
| rs190982 | 5 | 88,223,420 | $5.88 \times 10^{-01}$ | $3.2 \times 10^{-08}$ | - |
| rs1980493 | 6 | 32,363,215 | $7.83 \times 10^{-01}$ | - | $1.57 \times 10^{-08}$ |
| rs3129871 | 6 | 32,406,342 | $3.72 \times 10^{-01}$ | - | $3.43 \times 10^{-01}$ |
| rs3129882 | 6 | 32,409,530 | $1.04 \times 10^{-01}$ | - | $3.36 \times 10^{-02}$ |
| rs9268856 | 6 | 32,429,719 | $8.12 \times 10^{-01}$ | - | $5.51 \times 10^{-09}$ |
| rs9268877 | 6 | 32,431,147 | $1.91 \times 10^{-01}$ | - | $1.05 \times 10^{-08}$ |
| rs9271192 | 6 | 32,578,530 | $4.31 \times 10^{-01}$ | $2.9 \times 10^{-12}$ | - |
| rs10948363 | 6 | 47,487,762 | $8.62 \times 10^{-01}$ | $5.2 \times 10^{-11}$ | - |
| rs1020004 | 7 | 12,255,778 | $1.35 \times 10^{-03}$ | - | $4.59 \times 10^{-01}$ |
| rs6966915 | 7 | 12,265,988 | $1.24 \times 10^{-08}$ | - | $1.21 \times 10^{-01}$ |
| rs1990622 | 7 | 12,283,787 | $1.44 \times 10^{-08}$ | - | $7.88 \times 10^{-02}$ |
| rs2718058 | 7 | 37,841,534 | $9.99 \times 10^{-01}$ | $4.8 \times 10^{-09}$ | - |
| rs1476679 | 7 | 100,004,446 | $7.71 \times 10^{-01}$ | $5.6 \times 10^{-10}$ | - |
| rs11771145 | 7 | 143,110,762 | $9.67 \times 10^{-01}$ | $1.1 \times 10^{-13}$ | - |
| rs28834970 | 8 | 27,195,121 | $6.38 \times 10^{-02}$ | $7.4 \times 10^{-14}$ | - |
| rs3849942 | 9 | 27,543,281 | $8.94 \times 10^{-01}$ | - | $4.38 \times 10^{-04}$ |
| rs10838725 | 11 | 47,557,871 | $8.60 \times 10^{-02}$ | $1.1 \times 10^{-08}$ | - |
| rs10792832 | 11 | 85,867,875 | $4.50 \times 10^{-01}$ | $9.3 \times 10^{-26}$ | - |
| rs302668 | 11 | 87,876,911 | $4.57 \times 10^{-02}$ | - | $2.44 \times 10^{-07}$ |
| rs11218343 | 11 | 121,435,587 | $1.35 \times 10^{-01}$ | $9.7 \times 10^{-15}$ | - |
| rs17125944 | 14 | 53,400,629 | $1.71 \times 10^{-01}$ | $7.9 \times 10^{-09}$ | - |
| rs10498633 | 14 | 92,926,952 | $8.32 \times 10^{-01}$ | $5.5 \times 10^{-09}$ | - |
| rs242557 | 17 | 44,019,712 | $8.89 \times 10^{-01}$ | - | $4.82 \times 10^{-03}$ |
| rs8070723 | 17 | 44,081,064 | $7.48 \times 10^{-01}$ | - | $2.80 \times 10^{-04}$ |
| rs4147929 | 19 | 1,063,443 | $4.78 \times 10^{-01}$ | $1.1 \times 10^{-15}$ | - |
| rs2075650 | 19 | 45,395,619 | $2.11 \times 10^{-04}$ | - | $8.81 \times 10^{-07}$ |
| rs3865444 | 19 | 51,727,962 | $2.02 \times 10^{-01}$ | $3.0 \times 10^{-06}$ | - |
| rs7274581 | 20 | 55,018,260 | $3.01 \times 10^{-01}$ | $2.5 \times 10^{-08}$ | - |

**Figure 9.**
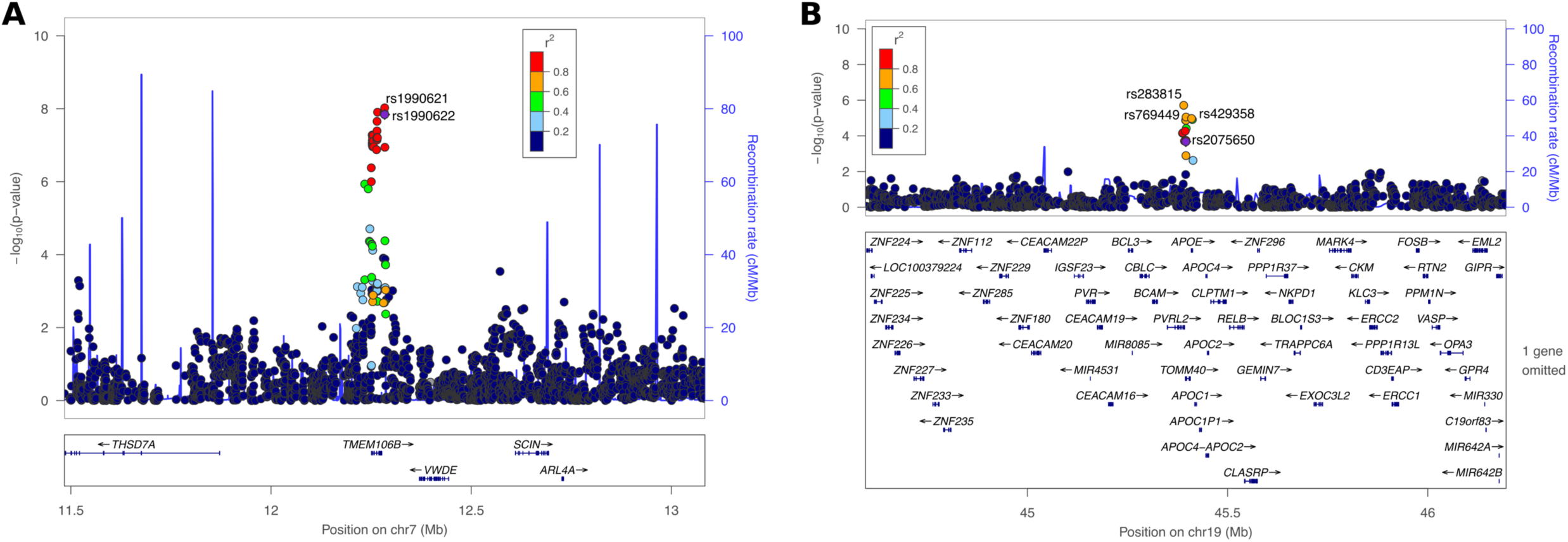
*TMEM106B* and *APOE* regions local plot. A) Local plot showed the zoom-in view of the hit in chromosome 7 with target SNP rs1990622 labeled with dark purple, and the top leading SNP is rs1990621. Nearby SNPs were also mainly located in the *TMEM106B* gene region and color coded with LD r2 thresholds. B) Local plot showed the zoom-in view of the hit in chromosome 19 with target SNP rs2075650 labeled with dark purple, and the top three leading SNPs are rs283815, rs769449, and rs429358. Nearby SNPs were also mainly located in the TOMM40/APOE gene region and color coded with LD r2 thresholds. One gene omitted in this region is SNRPD2.

In a gene-based analysis of our neuronal proportion QTL, *TMEM106B* (p-value = 2.96×10^−08^) is the only gene that passed genome-wide significant threshold followed by *APOE* (p-value = 3.2×10^−05^), the most important gene for sporadic AD risk (**Figure 6CD**). Previous GWAS for AD risk performed with the International Genomics of Alzheimer’s Project (IGAP) data stratified by *APOE* genotype showed that AD risk is significantly influenced by the interaction between *APOE* and *TMEM106B*^61^. Together with our observation of cellular composition QTL, these results suggest a potential interaction of *TMEM106B* and *APOE* may play a role in affecting AD risk/vulnerability and cellular composition balance between neurons and astrocytes, and the endosome and lysosome compartments might be the location that the interaction takes place.

## Discussion

The common variant rs1990622 in *TMEM106B* was first identified to be associated with FTD with TDP-43 inclusions^25^. Hyper-phosphorylated and ubiquitinated TDP-43 is the major pathological protein for FTD and ALS^62^, which is also present in a broader range of neurodegenerative disorders, including AD^63^, Lewy body disease^64^, and hippocampal sclerosis^63^. Recent study also suggested distinct TDP-43 types present in non-FTD brains, typical TDP-43 α-type and newly characterized β-type^65^. TDP-43 α-type is the typical form conventionally observed in temporal, frontal and brainstem regions. TDP-43 β-type is characterized by its close adjacency to neurofibrillary tangles, which is predominantly observed in limbic system, including amygdala, entorhinal cortex, and subiculum of the hippocampus. These findings suggested that pathologic TDP-43 protein that closely associated with *TMEM106B* variants might be the common pathologic substrate linking these neurodegenerative disorders. Multiple lines of evidence have merged and shown that protective variants in *TMEM106B* are associated with attenuated cognitive deficits or better cognitive performance in ALS^66^, hippocampal sclerosis^67^, presymptomatic FTD^68^, and aging groups with various neuropathological burden^69^ or in the absence of known brain disease^70^. Our study identified a protective variant rs1990621 of *TMEM106B* is associated with increased neuronal proportion in participants with neurodegenerative disorders and normal aging in non-demented controls. However, this effect is not observed in a younger schizophrenia cohort with a mean age of death less than 65 years old. This result suggested a common pathway involving *TMEM106B* shared by aging groups in the present or absence of neurodegenerative pathology that may contribute to cognitive preservation and neuronal protection.

Our study has demonstrated that a protective variant rs1990621 identified in *TMEM106B* gene region may exert neuronal protection function in aging groups. A protein coding variant rs3173615 in high LD with rs1990621 (r^2^ = 0.98) produces two protein isoforms (p.T185S). The S185 allele is protective and the protein carrying this amino acid is degraded faster than the risk variant T185. Thus, the risk allele of this coding variant leads to increased TMEM106B protein level^55,58,59^. *TMEM106B* overexpression results in enlarged lysosomes and lysosomal dysfunction^55,71^. It has also been shown that TMEM106B may interact with PGRN (the precursor protein for granulin) in lysosome^59^. Although rs3173615 is not included in our genomic data, it is in complete linkage disequilibrium with rs1990621 and rs1990622. It is worth pointing out that the minor allele of rs1990622, which has a protective effect in FTD, is in-phase with the minor allele of rs1990621, which is associated with increased neuronal proportion in our analysis. Despite the fact that our dataset is focused on neurodegeneration, we only have 11 verified FTD cases suggesting that TMEM106B might have a general neuronal protection role in neurodegeneration apart from FTD.

This observation suggested that a potential involvement of *TMEM106B* in the endosome/lysosome pathway may play a role in neurodegenerative disorder risk or vulnerability. Neuronal survival requires continuous lysosomal turnover of cellular contents through endocytosis and autophagy^72^. Impaired lysosomal function reduces lysosomal degradative efficiency, which leads to abnormal build-up of toxic components in the cell. Impaired lysosomal system has been found to be associated with a broad range of neurodegenerative disorders, including AD^73^, Parkinson disease^74-76^, Huntington disease^77,78^, FTD^79^, ALS^77^, Niemann-Pick disease type C^80,81^, Creutzfeldt-Jakob disease^82^, Charcot-Marie Tooth disease type 2B^83^, Neuronal ceroid lipofuscinoses (Batten disease)^84,85^, autosomal dominant hereditary spastic paraplegia^86^, Chediak-Higashi syndrome^87^, inclusion body myositis^88^, and osteopetrosis^89^. Considering the extensive involvements of lysosomal/endosomal compartments in neurodegenerative disorders, it has been proposed that a long and chronic process of abnormal metabolic changes during aging has led to the accumulation of toxic materials^72^. When lifespan increases especially in the sporadic forms of neurodegenerative disorders, failures to degrade these waste products break the proteostasis and the balance maintained by the multicellular interactions, and trigger subsequent chain reactions that lead to neuronal death and outbreaks of various neurodegenerative disorders due to different genetic susceptibilities and other disease etiologies. Although each neurodegenerative disorder has its own characteristic proteopathy, the boundaries of protein pathology distribution are never clear-cut across different disorders. In fact, copathology or nonspecific pathology of proteopathy have been observed in most autopsies of neurodegenerative disorders, such as TDP-43 discussed above, Lewy body, α-synuclein^90^, and etc. Our observation of lysosomal gene *TMEM106B* associated with neuronal proportion in aging cohorts suggests that the lysosomal pathway might be involved in the common mechanism underlying a broad range of neurodegenerative disorders or aging process in general that contribute to neuronal cell death.

Our study has demonstrated the great potential of using cell type composition as quantitative traits to identify QTLs associated with the changes in cell fractions. This approach is more powerful for disorders that involve considerably changes in cellular composition, for example, neurodegenerative disorders, and normal conditions during developmental or aging processes. The development of recent single cell studies will greatly increase the resolution in advancing our knowledge of cellular population changes. More detailed fine mapping of cellular composition from single cell studies together with machine learning algorithms, bulk RNA-Seq deconvolution will be more accurately capturing cellular fraction changes in the samples, such as different types of neurons or different states of astrocytes or microglia. Regarding scalability, this single cell powered bulk deconvolution approach is preferable for carrying out such cell type composition QTL analysis, because due to the high cost of performing single cell studies, bulk RNA-Seq is more financially feasible to scale up, and with larger sample sizes more hidden signals will be unrevealed with increased statistical power.

To conclude, we have identified a protective variant rs1990621 in *TMEM106B* associated with increased neuronal proportion through bulk RNA-Seq deconvolution and cell type proportion QTL analysis. This observation also replicated previous findings of the protective variant rs1990622 in FTD risk, which is in high LD with rs19990621^25^. Besides, we also observed the C allele of rs429358 (codetermine *APOE* ε4 isoform with rs7412 C allele) associated with decreased neuronal proportion as it was hypothesized. It suggested potential involvements of both *APOE* and *TMEM106B* in neuronal protection mechanisms underlying neurodegenerative and normal aging processes, and supported previous observation of interactions between these two genes^61^ in AD cohort. We speculate that *TMEM106B* related lysosomal changes might be involved in the common pathway underlying neuronal death and astrocytosis in neurodegenerative disorders and normal aging cohorts. With larger sample size and higher deconvolution resolution, this approach will reveal more biologically relevant and novel loci associated with changes in cellular composition that are important for understanding both disease etiology and healthy aging.

